# A mathematical model informs optimal fungicide use against Sclerotinia stem rot to maximize profits in soybean production

**DOI:** 10.1101/2024.10.30.621166

**Authors:** Tanner Byer, Tad Hatfield, Claus Kadelka

## Abstract

*Sclerotinia sclerotiorum*, the causative agent of stem rot (SR), is a significant yield-limiting disease affecting soybean crops in the temperate climates around the globe. Effective disease management practices rely on fungicides to mitigate the growth and spread of the disease. To infer optimal, profit-maximizing fungicide application rates, this study develops a mathematical model of mold and soybean growth with a requisite profit function. Sensitivity of the optimal fungicide application rate was computed against profit parameters (fungicide cost and soybean bushel price), and model parameters (mold growth rate, maximal SR damage to crops and fungicide efficiency). Expectantly, higher soybean bushel prices, rates of mold growth, and maximal mold damage to crops return elevated optimal fungicide rates. In contrast, higher levels of fungicide efficiency motivate lower optimal fungicide rates. The model also reveals a discontinuity in the optimal fungicide application rates for elevated fungicide costs; in this economic context, it becomes more profitable to apply no fungicide rather than low, ineffectual amounts that still allow mold to reach near-maximal outbreak levels in a finite time period. Future refinements of the model will incorporate variable mold growth rates modeled on annual weather patterns, crop rotation practices, and further exploring the relationships that soybean densities and row spacing have on mold growth, in order to build a more robust system to analyze the long-term effect of disease behavior on soybean crop yield.

## Introduction

Pathogenic species are rightly regarded as existential threats to human health through direct infection, however considerations to the proximal threat that pathogenic diseases impart on the scarcity of the world food supply are less common. A study conducted in 2021 by the Food and Agriculture Organization of the United Nations estimates that more than 220 billion U.S. dollars of crops are lost annually to pathogenic pests [1]. Particularly at risk is the most important oilseed crop worldwide (*Glycine max*), better known as the soybean. Annual global loss of soybean yield to pests and disease is estimated at 21.4% [2]. Integral to protecting soybean yield is the implementation of preventative measures against Stem rot (SR), a ubiquitous crop disease caused by the soil-borne fungal pathogen, *Sclerotinia sclerotiorum*. This pathogen affects large crop operations in temperate climates around the globe [3]. In North America alone, this yield-limiting fungus destroyed an average of 17 million bushels of soybean crops in 1996− 2009; economic estimates of cumulative profit loss surpassed one billion U.S. dollars for this period [4].

*S. sclerotiorum* propagates in the soil and can endure for three to eight years *in situ* [5]. After colonization of the fallen soybean petals, the fungus sprouts to release ascospores from structures known as apothecia [3]. A single sclerotium can produce more than two million ascospores in ten days; these ascospores disperse into the canopy of soybean crops and infect the nodes and stems of the surrounding crop [3, 6]. Consequently, if left untreated, SR outbreaks risk infecting crop operations beyond the bounds of a single farm operation. Epidemic disease dynamics are instigated by the formation of thick soybean canopies during the flowering stage of the plant life-cycle [7]. It is believed that the cool temperatures and moist soil conditions imparted by canopy formation catalyze the pathogenic behavior [4]. Consequently, the yearly incidence of SR fluctuates greatly in relation to annual temperature and rainfall levels, two factors associated substantially with disease dynamics. During seasons with low average temperatures and high precipitation, SR consistently ranks in the top five of the 23 most pernicious diseases that affect crop yield in North America [4].

To limit the detrimental effects of SR, both biotic and abiotic techniques are employed. Fungicide application constitutes one of the most prevalently used practices. Currently, chemical control of SR with foliar-applied fungicides is never 100% effective [8]. The amount of chemical control one can attain depends on the fungicide type, the application timing, and the spray coverage. In university trials, the amount of control from fungicides has ranged from 0 − 60% [4]. Boscalid and Picoxystrobin have been found to be the most effective fungicides [9]. The optimal timing of the fungicide application to control SR constitutes an active area of research [10]. It has been shown that fungicide application during the R1 growth stage provides a higher level of control than during the R3 stage of development [4]. Frequently, farmers apply fungicide multiple times in a growing season for improved disease control; considerations for application rates include residual fungicide duration and length of the soybean flowering stage [11]. Highly effective fungicides typically cost more, establishing a trade-off between fungicide effectiveness and cost [10]. Another area of research has focused on determining the efficacy of various fungicides in reducing the incidence of SR across different climates [12]. In drier years, when disease incidence is lower, one is less likely to see a return on fungicide investment than in wetter, high-incidence years [13]. Besides chemical control, SR is also assuaged by abiotic techniques including crop rotation (biennial soybean seasons), tillage, extended row spacing to limit elevated soil moisture, and lower seeding rates. [4, 14–16].

A number of approaches have been used to model SR growth [17–22]. Largely, these statistical models calculate probabilities of apothecial presence in the field as a function of hourly temperatures, humidity and wind speed [17, 20]. Other methods developed risk assessment algorithms to forecast disease severity in crop fields over seven day intervals based on canopy growth and soil micro-climate parameters [18, 19]. Machine learning models are also employed to predict incidence of disease from factors including prolonged leaf wetness and air temperatures [22]. Implementation of these methods requires frequent, precise, regio-specific data collection. However, many countries affected by the disease lack reliable data collection techniques, delaying coordinated responses to prevent disease establishment and spread [23]. Thus, mathematical models are needed to better understand the long-term outbreak behavior of the disease and thereby inform more efficient and enduring mitigation strategies to maximize crop yield and total farm revenue.

In this study, we develop a discrete model to optimize a static fungicide application rate to maximize profit from annually planted soybean crops over a multi-year period, given variation in the cost of fungicide and the market price for soybean bushels. Through a five-dimensional sensitivity analysis, we explore how the optimal rates are affected by changes in parameters related to mold growth, mold damage and fungicide efficiency. Although the complex, multistage disease life-cycle is markedly affected by stochastic elements not easily captured by mathematical models, we consider a simplified, abstract approach to capture the long-term behavior of the system and general qualitative effects.

## Methods

### Mold growth model

We describe the long-term growth of mold on a closed farm ecosystem using a modified logistic difference equation. Given the lack of detailed data regarding mold growth, we consider time in years. That is, within a single time unit multiple events take place:

1. Each year, *s* soybean seeds get planted per acre at the beginning of the season.
2. A proportion *β* of the planted seeds germinates and grows into plants, yielding a total of *βs* plants per acre. Previous work indicates approximate germination and emergence rates of 90% each [24, 25]. Thus, we set *β* = 81%.
3. The amount of mold in year *t* + 1 depends on the amount of mold in year *t*, with mold growing logistically at a maximal rate of *r* ∈ (0, 4). Since the canopy of soybean plants provides the requisite conditions for mold proliferation, the mold carrying capacity in the closed system is determined by the number of plants per acre scaled by a plant-to-mold conversion factor *κ*.
4. Fungicide application at a rate *ϕ* destroys a proportion *q* = *q*(*ϕ*) of all mold in a growing season. Previous work indicates diminishing returns at higher fungicide application rates [26, 27], implying

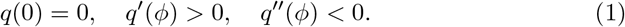

In the absence of specific data, we thus set *q*(*ϕ*) = log_*a*_(*ϕ* + 1) = ln(*ϕ* + 1)*/* ln(*a*), a relatively simple one-parameter function satisfying the requirements. As in [26], we assume that fungicide use beyond a certain point is detrimental to crop yield due to e.g. phytotoxicity or negative effects on beneficial organisms. We consider this point to be represented by *ϕ* = 1, which restricts reasonable fungicide use rates to *ϕ* ∈ [0, 1]. Since *q* describes a proportion, we also require *q*(1) ≤ 1. This implies the meaningful range for the base of the logarithm is *a* ∈ [2, ∞). Note that the value *q*(1) = log_*a*_(2) describes the maximal fungicide-induced proportion of mold that can be destroyed by fungicide (if applied at rate *ϕ* = 1) over the course of one season (Fig. 1A). We use this parameter as a proxy for the efficiency of the fungicide in use. By default, we set *a* = *e* ≈ 2.718 such that *q*(*ϕ*) = ln(*ϕ* + 1).

**Fig 1.**
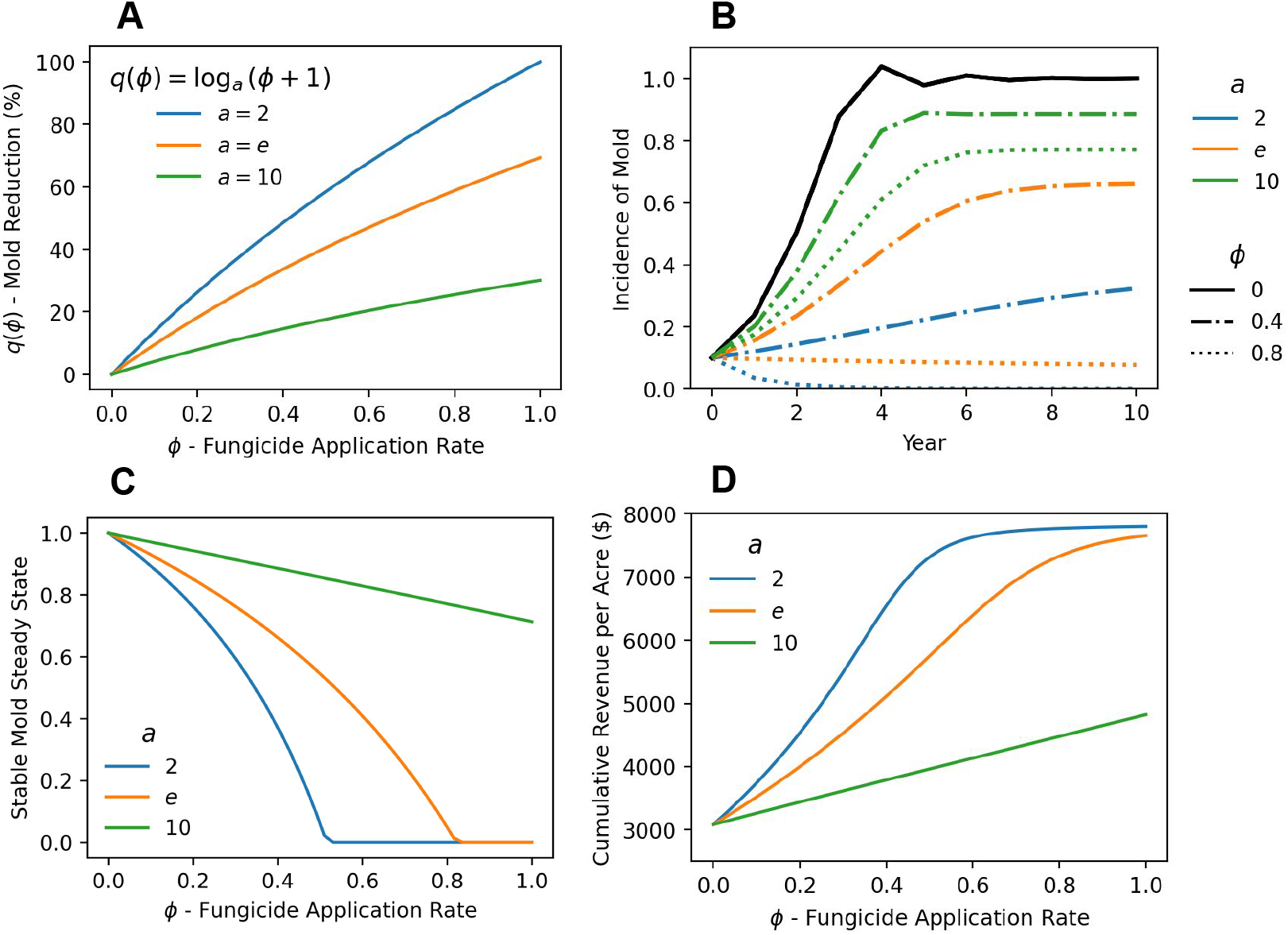
Fungicide efficiency and application rate impacts mold growth and maximal cumulative soybean profits. (A) The fungicide-induced mold reduction (y-axis) depends logistically on the fungicide application rate *ϕ* (x-axis) and the fungicide efficiency (scaled by *a*, colors). (B) Mold incidence over ten years assuming a constant fungicide application rate *ϕ* (line styles) and fungicide efficiency (scaled by *a*, colors). Other mold growth parameters are set at the default values, specified in Table 1. (C) Stable mold steady states for variable levels of fungicide application and efficiency. Mold growth parameters were chosen at default values (Table 1). When *r*(1 − *q*(*ϕ*)) *<* 1, the zero steady state is stable. (D) Cumulative revenue (Eq. 6) over ten years. Mold growth and revenue parameters are at their default values (Tables 1, 2).

The following equation describes the change in mold per acre from one year to the next:

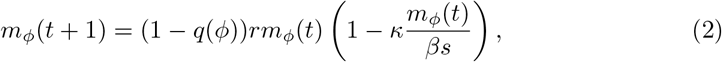

By considering a constant intrinsic mold growth rate and constant germination rate, we implicitly assume that the soil moisture and temperature are consistent across years. Given the year-to-year variability in these values, a stochastic model would constitute a better but also more complicated representation of reality.

In the absence of fungicide, the non-trivial mold outbreak steady state is

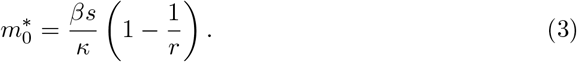

In the context of our model, this is the highest level of mold the farm can experience given a fixed mold growth rate and a fixed number of seeds planted and germinating each year. By setting

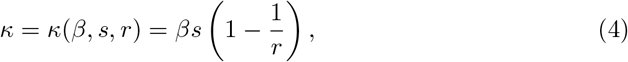

we have 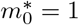. This choice of *κ* thus enables us to interpret *m*_*ϕ*_(*t*) as a proportion of the maximally possible long-term mold coverage. For any fungicide application rate *ϕ ≥* 0, the non-trivial mold outbreak steady state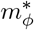 can be derived from Eq. 2 and Eq. 4.

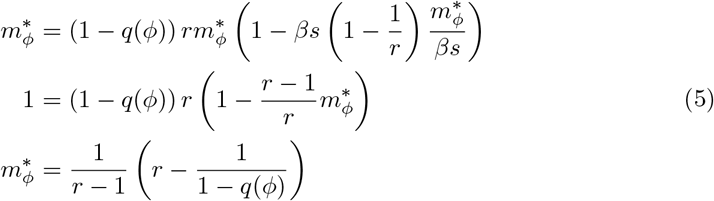

Since *q*(*ϕ*) > 0 whenever *ϕ* > 0 (Eq. 1), we have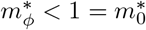.

We choose the following default parameters: *s* = 140, 000 seeds/acre, *β* = 81% of seeds germinating and turning into soybean plants, a maximal mold growth rate of *r* = 2.5, and *m*_*ϕ*_(0) = 10% of the farm is initially covered by mold (see Table 1). The fungicide application rate is varied on the interval [0, 1] as part of the optimal control problem defined below.

**Table 1.**
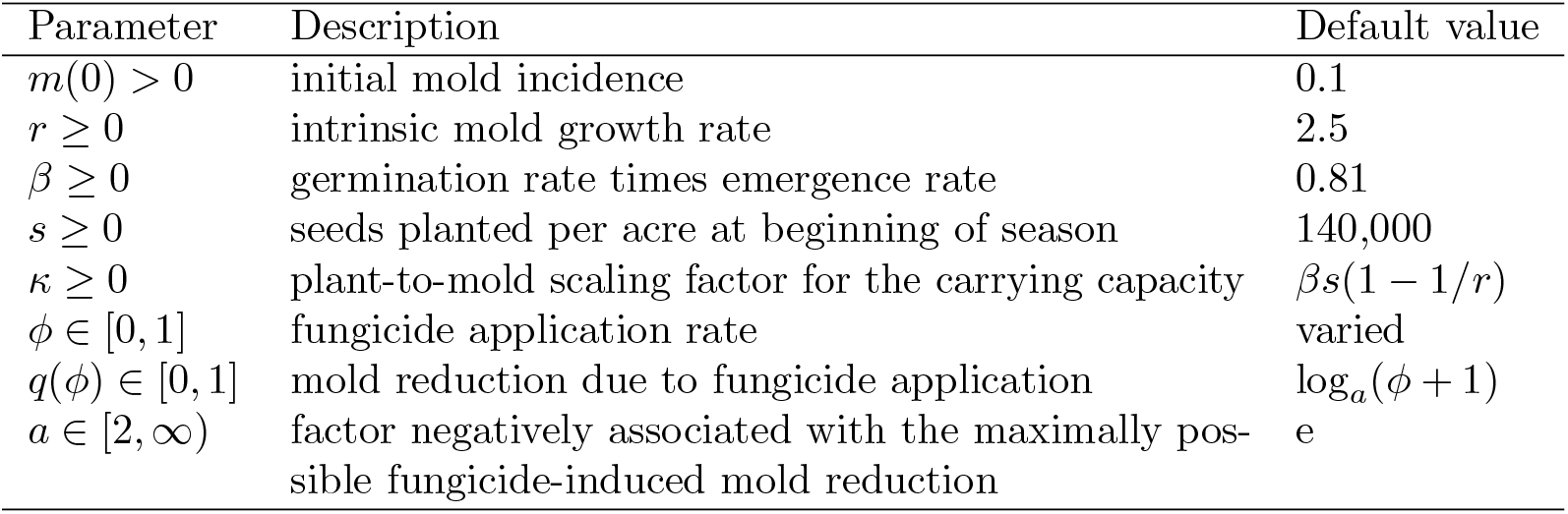
Model parameters.

| Parameter | Description | Default value |
| --- | --- | --- |
| $m(0) > 0$ | initial mold incidence | 0.1 |
| $r \geq 0$ | intrinsic mold growth rate | 2.5 |
| $\beta \geq 0$ | germination rate times emergence rate | 0.81 |
| $s \geq 0$ | seeds planted per acre at beginning of season | 140,000 |
| $\kappa \geq 0$ | plant-to-mold scaling factor for the carrying capacity | $\beta s(1 - 1/r)$ |
| $\phi \in [0, 1]$ | fungicide application rate | varied |
| $q(\phi) \in [0, 1]$ | mold reduction due to fungicide application | $\log_a(\phi + 1)$ |
| $a \in [2, \infty)$ | factor negatively associated with the maximally possible fungicide-induced mold reduction | $e$ |

### Optimal mold control

The seasonal soybean revenue is the product of the total soybean bushels produced (*ψβs*) and the average market price of a soybean bushel (*b*), where *ψ* describes the plant to bushel conversion rate. Recent data from Eastern South Dakota indicates that *s* = 140, 000 seeds per acre on average result in *ψβs* = 65 bushels of soybeans [28]. With *β* = 0.81, this yields *ψ* = 1*/*1744. We acknowledge that soybean yield depends on many factors besides the initial number of seeds and germination rates [29]. Some of these factors are not accounted for in the model to keep it relatively simple and interpretable.

To model the effect of mold on the soybeans, we assume that only a proportion of the crops can be sold. If the mold incidence *m*_*ϕ*_(*t*) is equal to the worst possible and long-term mold incidence at no fungicide use,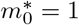 (Eq. 3), we assume that only 1 − *θ* of the crops can be sold. That is, *θ* describes the maximal damage of mold to plants, and we set *θ* = 70% as the default [30]. If the mold incidence *m*_*ϕ*_(*t*) is lower, mold reduces the yield at a proportionately lower rate.

Finally, the cost of fungicide application can be assumed to depend linearly on the number of plants on the farm. If *c* represents the cost of fungicide application per soybean plant, we thus have the following per-acre soybean profit function:

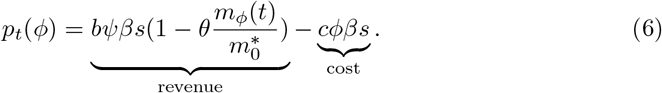

The one-dimensional control problem investigated here consists of choosing the optimal fungicide application rate *ϕ*\* ∈ [0, 1] that maximizes the profit over a fixed 10-year planning horizon. For simplicity, we do not discount future profits. That is,

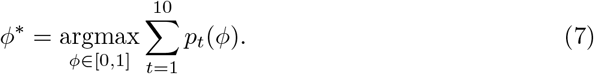

## Results

Given the lack of mathematical models to guide the management of SR on soybean farms, we developed a relatively simple difference equation model that describes the incidence of mold in a given season as a function of the mold surviving from the year prior and the mold-reducing fungicide application rate *ϕ*. Clearly, more fungicide use results in higher revenue but also higher cost. Given this trade-off, we investigated which constant rate of fungicide application maximized the per-acre profit over ten years.

Assuming logistic mold growth, standard knowledge of the logistic map (see e.g., [32]) implies that the mold incidence, modeled by Eq. 2, transitions to a stable steady state as long as the effective (fungicide use-adjusted) mold growth rate *r*_eff_ := *r*(1 − *q*(*ϕ*)) ∈ [0, 3) (Fig. 1B). Waning fungicide efficiency portends higher, stable steady states of mold for comparable fungicide application rates. At *r*_eff_ = 1, a transcritical bifurcation occurs: As *r*_eff_ increases, the zero steady state becomes unstable, while another steady state becomes positive and stable. As *r*_eff_ approaches 3, convergence to this steady state slows down. At *r*_eff_ = 3, a first period-doubling bifurcation occurs and a stable two-cycle emerges, followed by the emergence of a stable four-cycle, eight-cycle, etc. Therefore, a mold growth rate of *r* = 2.5 (the default choice here in the absence of reliable data) gives rise to stable long-term behavior for any choice of fungicide application rate *ϕ* ∈ [0, 1]. At low fungicide application rates, as long as *r*(1 − *q*(*ϕ*)) > 1, the mold will persist and transition towards the positive stable steady state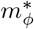 (Eq. 5). On the other hand, at high fungicide application rates (whenever *r*(1 − *q*(*ϕ*)) *<* 1), mold incidence will decrease logistically towards the stable zero steady state (Fig. 1C). In this case, 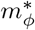 is an unstable negative steady state of Eq. 2.

### Partial model validation

One-dimensional parameter sensitivity analyses revealed mostly expected behavior. The annual profit (Eq. 6) increases if the soybean bushel price (*b*), the plant to bushel conversion rate (*ψ*), the number of seeds (*s*), the germination and emergence rate (*β*) and the maximal possible fungicide-induced mold reduction (log_*a*_(2)) are higher (Fig. 1D). Similarly, it follows directly from Eq. 6 that the annual profit decreases if the fungicide cost (*c*), the maximal mold damage (*θ*), the intrinsic mold growth rate (*r*), or the initial mold incidence (*m*(0)) are higher. This expected behavior partially validates the model. The fungicide application rate *ϕ* ∈ [0, 1] constitutes the only model parameter that has a non-trivial, non-monotonic influence on the profit (since an increase in *ϕ* increases both revenue and cost). The remainder of the manuscript explores the optimal choice of *ϕ* under a variety of conditions. Given the lack of reliable estimates for a number of key parameters, the investigated model is somewhat abstract and the focus is on qualitative, rather than quantitative findings.

### Optimal fungicide application rates

To infer the sensitivity of the optimal fungicide application rate *ϕ*\* ∈ [0, 1] to changes in soybean bushel price (*b*) and fungicide cost (*c*), we varied these two parameters, while fixing all other parameters at the default values described in Tables 1 and 2. At a fixed rate of fungicide application, the ten-year profit increases monotonically as the price per bushel increases (Fig. 2A-C) and as fungicide costs decrease (Fig. 2D-F). At the same time, the optimal fungicide application rate *ϕ*\* increases as the bushel price increases and decreases as the fungicide cost increases. We interpret these expected results as follows: when farmers earn more per soybean, there is an increased incentive to produce a higher yield through intensified fungicide application. The sensitivity of *ϕ*\* to changes in bushel prices decreases as the price increases. For very low bushel prices, the optimal fungicide application rate is, as expected, zero.

**Table 2.**
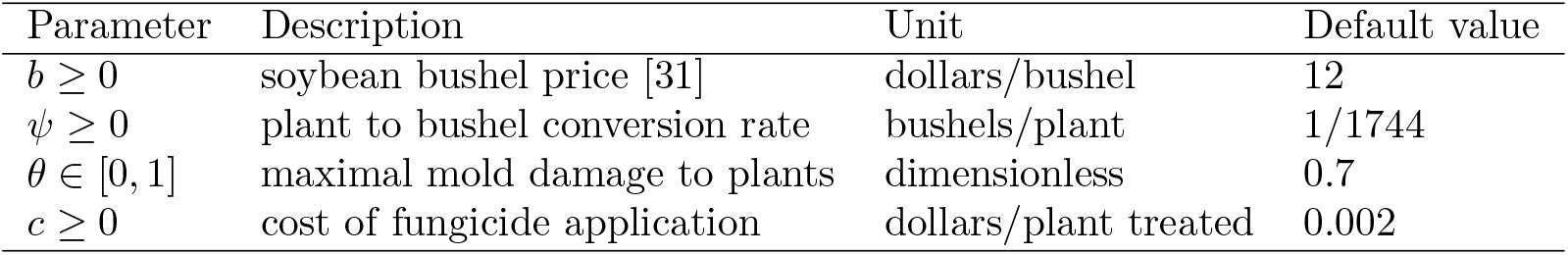
Profit function parameters.

| Parameter | Description | Unit | Default value |
| --- | --- | --- | --- |
| $b \geq 0$ | soybean bushel price [31] | dollars/bushel | 12 |
| $\psi \geq 0$ | plant to bushel conversion rate | bushels/plant | 1/1744 |
| $\theta \in [0, 1]$ | maximal mold damage to plants | dimensionless | 0.7 |
| $c \geq 0$ | cost of fungicide application | dollars/plant treated | 0.002 |

**Fig 2.**
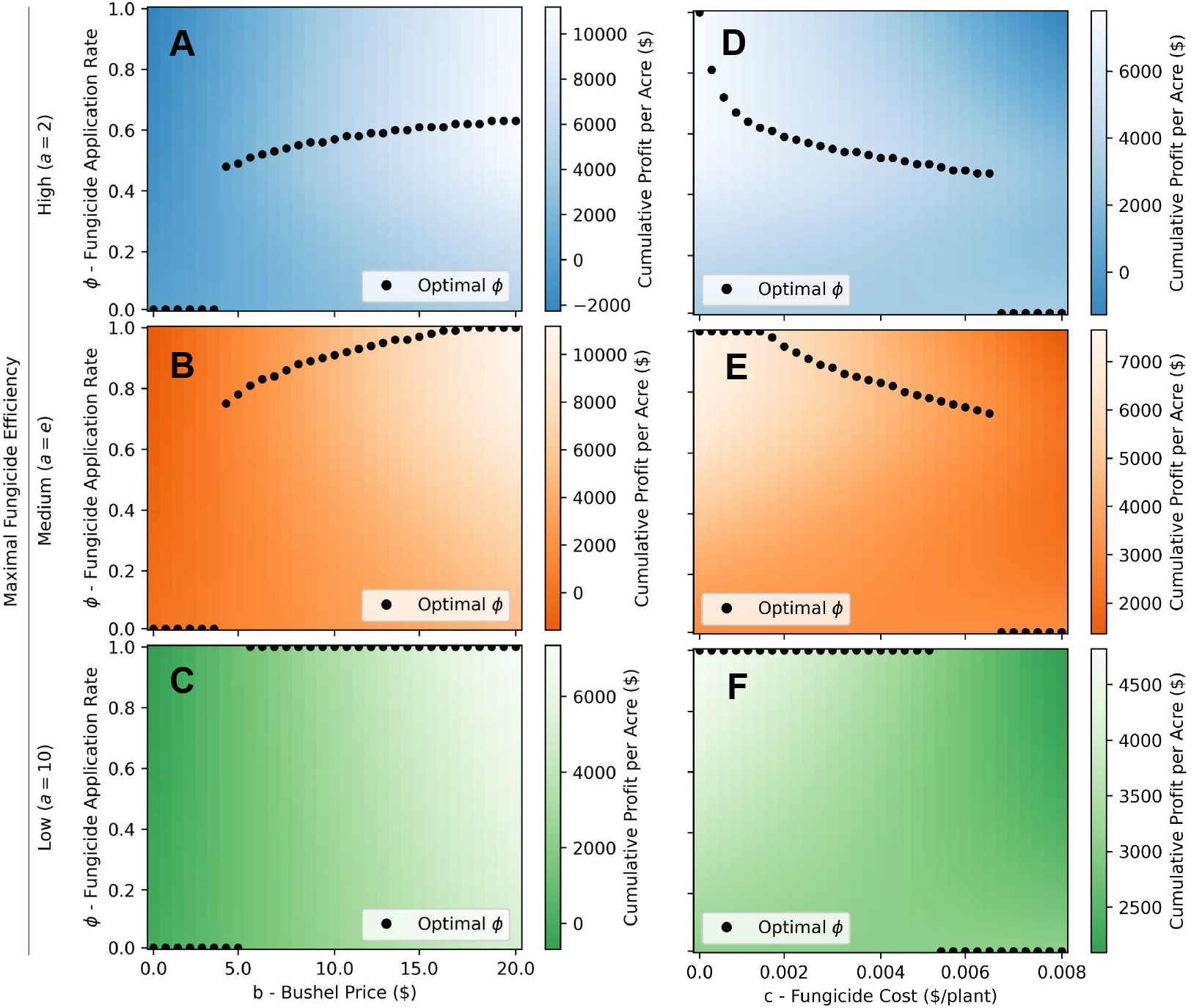
Sensitivity of the profit-maximizing fungicide application rate to soybean bushel price, fungicide cost, and fungicide efficiency. The cumulative ten-year profit per acre of soybeans is shown for different combinations of constant fungicide application rates *ϕ* ∈ [0, 1] (y-axis), and (A, B, C) soybean bushel prices *b* ∈ [0, 20] and (D, E, F) fungicide costs *c* ∈ [0, 0.006] (x-axis). The fungicide efficiency is (A,D) high (*a* = 2), (B,E) medium (*a* = *e*), and (C, F) low (*a* = 10). For all other parameters, default values are used (Tables 1, 2). Black circles indicate the optimal (i.e., profit-maximizing) constant fungicide application rate.

When the application of fungicide does not diminish the profit at all, farmers are encouraged to use it liberally to maximally offset any mold growth. This explains why *ϕ*\* = 1 if the cost *c* = 0 (Fig. 2D-F). Even at very low fungicide costs, *ϕ*\* quickly drops to more conservative rates that still prevent rapid mold outbreaks as long as fungicide efficiency is high (Fig. 2D). In these situations, mold is present and may even slowly grow but the incidence remains much lower over the ten-year time frame than 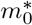, which is reached in the long-term absence of fungicide. On the other hand, if fungicide is less efficient (Fig. 2E,F), more plants need to be treated with fungicide to achieve the same reduction in mold growth, explaining the slower reduction in *ϕ*\* (as the cost increases) in these scenarios.

Interestingly, the optimal fungicide application rate as a function of bushel price and fungicide cost is not continuous. There exists a threshold where the optimal rate suddenly jumps (Fig. 2). A five-dimensional sensitivity analysis, in which both bushel price and fungicide cost are varied at the same time, reveals that this threshold is a linear function in these two parameters (Fig. 3). Moreover, the phenomenon is evident at all investigated levels of fungicide efficiency, mold damage to crops, and mold growth rates. If the maximal damage mold can inflict on crops is smaller (*θ* = 0.3 vs *θ* = 0.7) or if mold grows slower (*r* = 2 vs *r* = 2.5), the threshold bushel price, at which the use of fungicide becomes profit-maximizing, is higher. The size of the jump in optimal fungicide application rates increases as the fungicide efficiency decreases (Figs. 2,3). If the fungicide efficiency is low (*a* = 10), the optimal fungicide application rate takes on only extreme values: apply fungicide to all plants or none at all.

**Fig 3.**
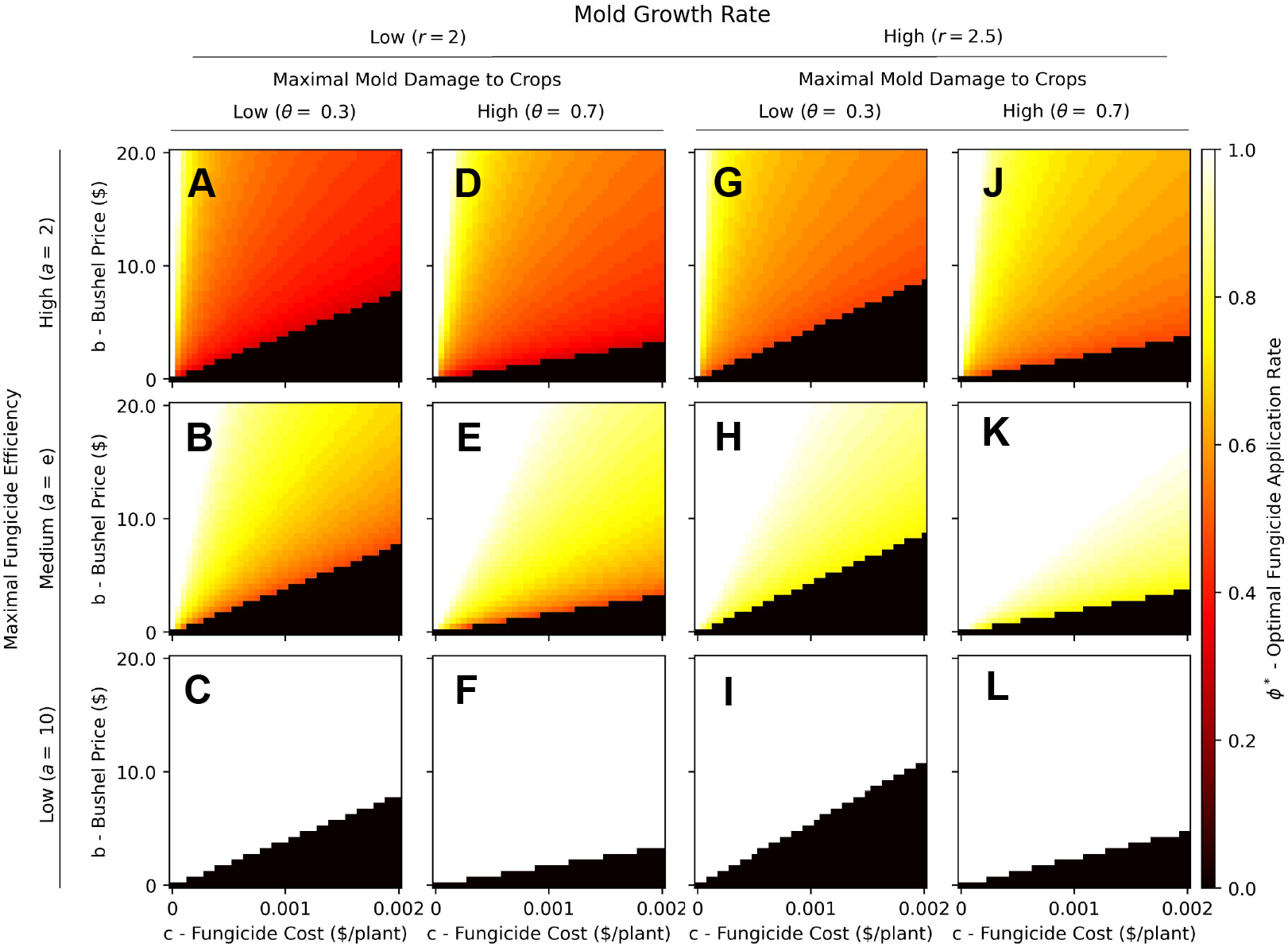
Five-dimensional sensitivity analysis of the profit-maximizing fungicide application rate. The optimal (i.e., profit-maximizing) fungicide application rate, *ϕ*\* ∈ [0, 1], (cell color) is shown for soybean bushel prices *b* ∈ [0, 20] (y-axis) and fungicide costs *c* ∈ [0, 0.002] (x-axis). Three further parameters are varied between the sub panels: (i) fungicide efficiency modulated by the parameter *a* ∈ {2 (high; first row), *e* (medium; second row), 10 (high; third row)}, (ii) maximal mold damage to crops, *θ* ∈ {0.3 (low; first and third column), 0.7 (high; second and fourth column)}, and (iii) mold growth rate *r* ∈{2 (low; first and second column), 2.5 (high; third and fourth column)}. Default values, specified in Tables 1,2), were used for all other parameters.

To understand reasons behind the occurrence of the jump discontinuity in optimal fungicide application rates, we considered the profit as a function of fungicide application rate *ϕ* for costs below, at and above the threshold where use of fungicide all of sudden becomes optimal (Fig. 4). Irrespective of the fungicide cost and efficiency, the function has two local maxima: one at *ϕ*_1_ = 0 corresponding to no fungicide use, and another at *ϕ*_2_ > 0. The optimal fungicide application rate is *ϕ*\* = max(*ϕ*_1_, *ϕ*_2_). As the fungicide cost decreases, eventually *ϕ*\* = *ϕ*_2_. On the other hand, when fungicide costs are sufficiently high, ineffectual fungicide application rates that allow mold growth to near outbreak steady state levels by year ten are less profitable than no fungicide application. The more efficient the fungicide, the smaller is *ϕ*_2_ because the amount of fungicide needed to effectively control mold growth (e.g., to achieve *r*_eff_ = *r*(1 − *q*(*ϕ*)) *<* 1) is lower. If fungicide efficiency is low (e.g., a=10, Fig. 4C), *ϕ*_2_ = 1 corresponding to applying the maximally feasible amount of fungicide. This explains the exclusive occurrence of extreme values *ϕ*\* ∈ {0%, 100%} in this scenario (Fig. 2C,F, and Fig. 3C,F,I,L).

**Fig 4.**
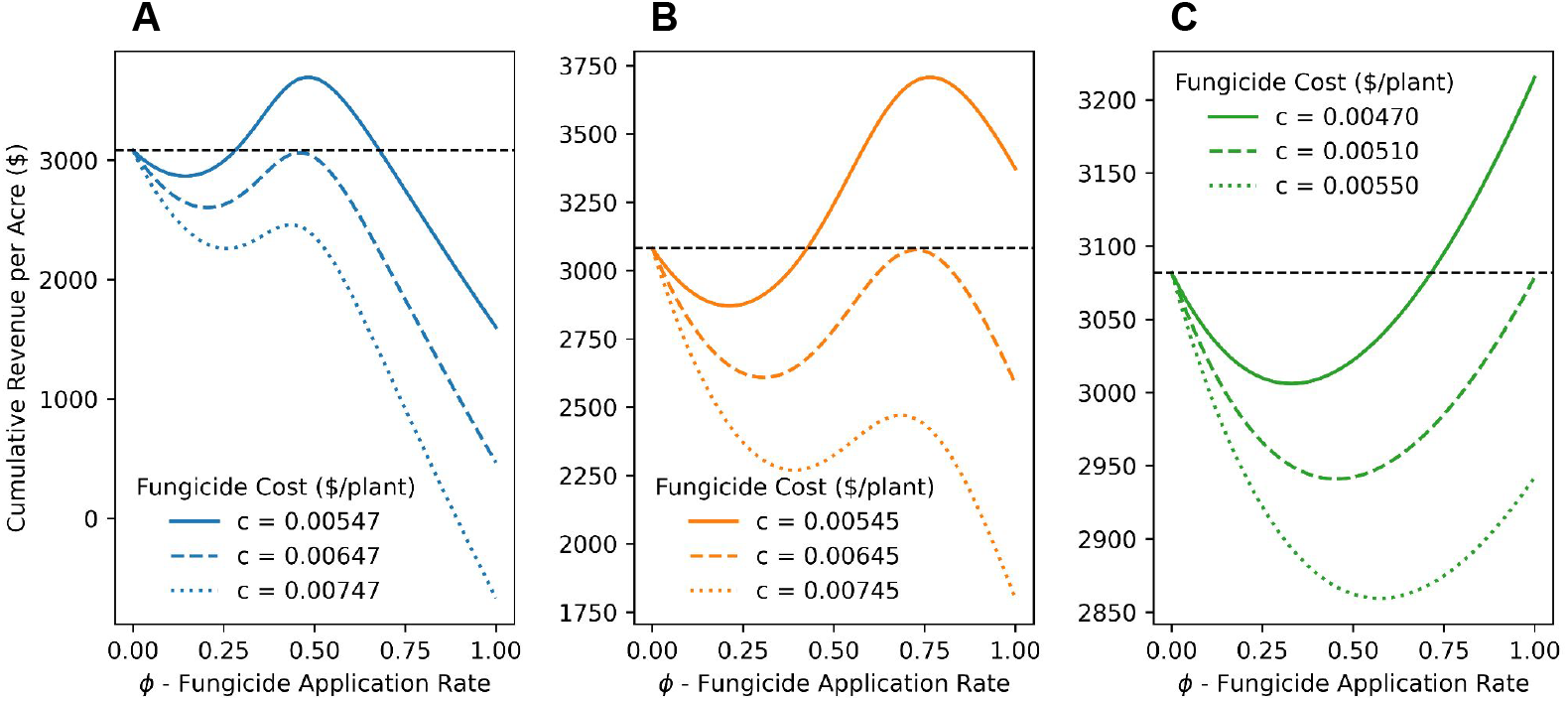
Profit as a function of fungicide application rate for different costs. The cumulative ten-year profit per acre (y-axis, Eq. 6) is plotted against the fungicide application rate (*ϕ* ∈ [0, 1], x-axis), for three specified fungicide costs *c*, and three levels of fungicide efficiency: (A) high (*a* = 2), (B) medium (*a* = *e*), (C) low (*a* = 10). Default values from Tables 1,2 are used for all other parameters.

## Discussion

In this study, we developed a logistic growth model for soybean-affecting stem rot and a soybean profit function to discern the optimal fungicide application rates to maximize the cumulative soybean profit over a finite ten-year period. Sensitivity analyses of model and profit parameters revealed mostly expected effects: increased optimal rates of fungicide use when soybean prices rise, fungicide costs fall, and mold growth rates and maximal mold-induced damage to soybean plants are elevated. The model analysis also revealed a highly sensitive parameter region where small perturbations in soybean bushel price or fungicide cost induce large changes in calculated optimal fungicide application rates. We showed that this jump discontinuity occurs when applying an ineffectual amount of fungicide allows the disease to reach near-maximal levels, which is less profitable than purchasing and applying no fungicide. In light of the increased effectiveness of fungicides on limiting mold growth, these results suggest a more tempered approach to fungicide application to ensure maximal profit [9].

The lack of detailed data to estimate parameters related to mold growth, fungicide efficiency, and maximal mold damage to soybean yield greatly limits our modeling efforts. This explains why we developed a relatively simple discrete-time model and focused on a qualitative (rather than quantitative) interpretation of the results. For example, we note that the specific jump discontinuity in optimal fungicide use depends on parameter values (Fig. 3), but rather than ascertain parameter values at the discontinuity, we focus on explaining why this region exists.

The observed high sensitivity of optimal fungicide application rates to changes in macroeconomic and ecological parameters highlights the importance of continued investigation of the factors underlying mold growth and damage to soybean plants. Recent advances in data gathering techniques that employ remote sensing technologies have enhanced our understanding of real-time SR levels and their impact on crops [33]. Such data are crucial for refining predictive models. For instance, Sporecaster, a collaborative effort led by the University of Wisconsin, leverages statistical modeling within a mobile app to assess in-season risks of SR [34]. Improving mathematical models by incorporating real-time data will help build a better framework of tools to be used by the agricultural community.

Due to the lack of robust modeling data, we built a simple logistical model and profit function. Future access to more data would enable the formulation of a more complex model that accounts for stochastic factors that influence mold growth. More specifically, we do not account for abiotic variables and established practices such as crop rotation, planting different varieties of the same crop, weed coverage, field topography, and the influx of ascospores from outside fields. Further, we assume a constant mold growth rate and seed germination and emergence rate, even though these rates almost certainly depend on weather conditions. Our study is also limited by the inherent simplification of modeling mold growth in annual time steps. Given the complexity of the mold life cycle [4], we have substantially simplified the number of factors that contribute to the growth of mold to better understand the long-term behavior of the ecosystem.

Our model is further built on several assumptions that may not always hold true. Although the fungus can persist in the field for many years, the levels of the pathogen and the severity of the disease do not necessarily always rise with the continuous cultivation of susceptible crops [14, 35, 36]. Another assumption is the connection of disease incidence with yield; although there is a negative relationship between the two, this is not always clearly defined, especially at low incidence levels [8, 9, 37]

We note two further limitations. First, the predicted per-acre profit is much higher than realistic because we only account for costs related to fungicide use and not costs of labor, land rent, or repairs, which often exceed fungicide costs [38]. Nevertheless, the model results should still hold because some of these costs are fixed costs while others can be considered as part of the fungicide costs, e.g., higher labor costs associated with applying the fungicide. Second, we consider only a fixed amount of soybean seeds planted per acre, *s* = 140, 000, a number that has been shown to result in optimal yields [39]. However, the amount of seeds planted is known to affect the size of plants and the canopy (due to, e.g., resource competition due to plant overcrowding), and it may affect the spread of mold. Similarly, exploring a density-dependent relationship between the number of plants and the total crop yield that reflects the limitations of nutrients in the soil is another area of interest. From this exploration, we would be interested in optimizing both fungicide application rates and the number of seeds planted in a season to achieve maximum profit. At the same time, it may prove optimal to alter the level of fungicide use between years, something we do not explore in this initial analysis as (i) it would substantially complicate the optimal control problem and (ii)a practical implementation would require detailed measurements.

Lastly, we acknowledge that the developed model is general enough to represent, to variable degrees of accuracy, multiple fungus-plant relationships. Although *Sclerotinia sclerotium*, the causative agent of SR, is most widely known for its effects on soybeans, it also affects many different plants [23, 37]. Further, other fungal pathogens like *Claviceps purpurea* could also be modeled using our framework.

Altogether, the presented results are a first attempt to qualify the decisions faced by soybean farmers every season across large parts of North America: should we use yield-improving fungicide against SR, and if so, how much? It is important to regard the results presented here in a qualitative way to avoid a direct interpretation as we still lack detailed data for several key model parameters, and our model lacks the stochastic nature required to model real-world conditions.

## Author Contributions

Conceptualization: TB, TH, CK; Formal analysis: TB, TH, CK; Methodology: TB, TH, CK; Software: TB; Visualization: TB, CK; Supervision: CK; Writing: TB, TH, CK.

## Data and Materials Availability

The Python code used to run developed model and generate all figures is provided at https://github.com/ckadelka/mold-growth.

